# Utilizing ExAC to Assess the Hidden Contribution of Variants of Unknown Significance to Sanfilippo Type B Incidence

**DOI:** 10.1101/253435

**Authors:** Wyatt T. Clark, G. Karen Yu, Mika Aoyagi-Scharber, Jonathan H. LeBowitz

## Abstract

Given the large and expanding quantity of publicly available sequencing data, it should be possible to extract incidence information for monogenic diseases from allele frequencies, provided one knows which mutations are causal. We tested this idea on a rare, monogenic, lysosomal storage disorder, Sanfilippo Type B (Mucopolysaccharidosis type IIIB).

Sanfilippo Type B is caused by mutations in the gene encoding α-N-acetylglucosaminidase (NAGLU). There were 189 NAGLU missense variants found in the ExAC dataset that comprises roughly 60,000 individual exomes. Only 24 of the 189 missense variants were known to be pathogenic; the remaining 165 variants were of unknown significance (VUS), and their potential contribution to disease is unknown.

To address this problem, we measured enzymatic activities of 164 NAGLU missense VUS in the ExAC dataset and developed a statistical framework for estimating disease incidence with associated confidence intervals. We found that 25% of VUS decreased the activity of NAGLU to levels consistent with Sanfilippo Type B pathogenic alleles. We found that a substantial fraction of Sanfilippo Type B incidence (67%) could be accounted for by novel mutations not previously identified in patients, illustrating the utility of combining functional activity data for VUS with population-wide allele frequency data in estimating disease incidence.

## 1 Introduction

Accurate epidemiological data on rare disorders are extremely valuable, but collecting such information can be a difficult task for several reasons. Retrospective studies can often be inaccurate due to registries being incomplete or simply unavailable. Active screening can be more accurate, but is also more costly. Generating accurate estimates by screening for a rare disease requires coverage of a large number of births over a long period of time. Furthermore, while newborn screening is performed for several rare disorders; it is generally limited to cases where detection at birth is possible, and to those disorders where a method of intervention is available [1]. Such factors make it very difficult to estimate the incidence, or prevalence, of rare monogenic disorders that are either under-studied or that have no options for treatment.

A complementary method of estimating disease incidence would be to calculate the combined carrier rate of all mutations that contribute to pathology [2–4]. For a population in equilibrium one would simply need to know which variants cause disease, and obtain an estimate of their frequencies in the population of interest [5–7]. However, large scale sequencing data-sets contain an abundance of variants of unknown significance (VUS) which can be located in genes associated with monogenic disorders [8]. Although the allele frequency of individual VUS may be quite low, the combined allele frequency of all VUS in a given gene can be significant. Not taking such variants into consideration can potentially lead to the underestimation of disease incidence.

In order to determine the hidden contribution of VUS to disease incidence we used Mucopolysaccharidosis type IIIB (Sanfilippo Type B), caused by defects in the gene encoding α-N-acetylglucosaminidase (NAGLU), as a case study. We experimentally determined the residual enzyme activity associated with the NAGLU VUS found in ExAC using an *in vitro* cell-based assay. We found that a large number of NAGLU VUS had enzymatic activity levels low enough to be consistent with known Sanfilippo Type B pathogenic alleles. We showed that such variants make a significant impact on the allele frequency-based estimate of Sanfilippo Type B incidence, bringing it in line with literature reports.

We also describe a statistical framework for calculating confidence intervals associated with incidence estimates. This model accounts for differing sample sizes associated with each variant; an important feature when sample sizes can span orders of magnitude from one mutation to the next within the same dataset. We validated our method and found very little deviation between the confidence intervals generated using this framework and what was observed through bootstrapping simulation.

Finally, we show that resources such as ExAC present a large enough sample of the population to estimate disease incidence for disorders as rare as Sanfilippo Type B with acceptable error margins.

## 2 Results

### 2.1 The potential contribution of VUS to disease incidence can be significant

Figure 1 shows that VUS are likely to contribute significantly to the disease incidence of several lysosomal storage disorders. We compared literature-reported incidence rates with incidence rates calculated using pathogenic variants (Disease causing (DM) variants in HGMD) and loss of function (LOF) variants as defined in Section 4.3 found in ExAC in the Non-Finnish European population (Section 4.5, Methods). While the combination of previously identified disease mutations from HGMD and high confidence LOF variants made significant contributions to incidence estimates, for almost all disorders we observed that only considering these two categories of variants greatly underestimated disease incidence rates. For example, for Sanfilippo Type B we arrived at an estimated disease incidence of 1 in 1,091,549 using the allele frequencies of HGMD and LOF variants whereas epidemiological data estimate it to be 1 in 321,128 [9]. While there are several factors that might have an impact, the discrepancy between what is reported in the literature, we set out to test the contribution of VUS to the observed discrepancy.

**Fig 1.**
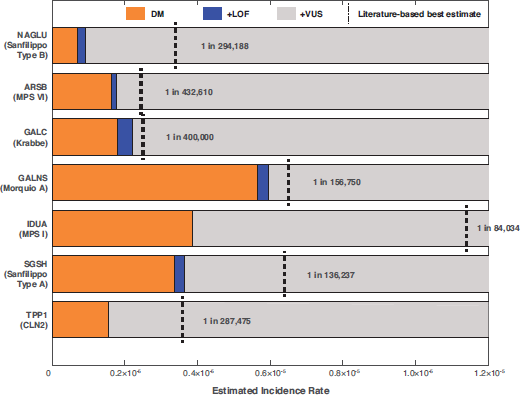
Allele frequency based incidence estimates of lysosomal storage disorders in the ExAC Non-Finnish Population. Starting with pathogenic (DM) variants in HGMD (orange), incidence was estimated using allele frequencies for each successive class of variant, combining mutations with all previous categories. Loss of function (LoF) variants (blue) were selected as those mutations causing either a splice affecting, stop-gained or frameshift change in the coding sequence which had not been documented in HGMD. Likewise, variants of unknown significance (VUS) were selected as those missense mutations in each gene that had yet to be documented in HGMD. Vertical dashed black lines represent the reported incidence rate of each disorder in the literature for the European region for Sanfilippo Type B [9], MPS VI [10], Krabbe [11], Morquio A (average of the UK, Germany, and the Netherlands) [12], MPS I [13], MPS IIIA [9], and CLN2 (average of Sweden, Norway, Finland, Italy, Portugal, Netherlands, the Czech Republic, and Italy) [2].

### 2.2 Many NAGLU VUS in ExAC have reduced enzymatic activity

In order to understand the contribution of VUS to disease incidence we developed an *in vitro* cell-based enzymatic activity assay to determine the functional impact of missense VUS in NAGLU. Individual cDNA’s for each NAGLU missense mutant were made and transfected in HEK293 cells. NAGLU enzyme activity was measured using a fluorogenic 2/21 assay with the substrate 4-Methylumbelliferyl-N-acetyl-α-D-glucosaminide (see Section 4.2). We tested 164 missense VUS found in ExAC v0.3. We also tested 35 deleterious missense mutations reported in HGMD, 17 of which were found in ExAC as well. As shown in Figure 2A, a range of enzymatic activities were observed in these NAGLU VUS. 41 out of 164 assayed VUS displayed enzymatic activities that were less than 15% of wild type (WT) NAGLU activity.

**Fig 2.**
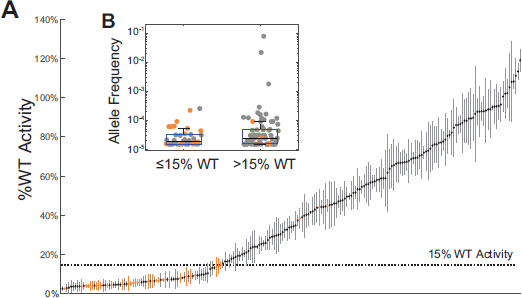
The enzymatic activity of NAGLU variants. **(A)** Variants are ordered by average %WT activity. Standard deviation in %WT activity in replicates is represented by vertical bars. Previously identified disease variants are shown in orange. A dashed line shows the 15%WT activity threshold below which variants are considered to be pathogenic. **(Insert B)** A box plot (y-axis log scale) of the Non-Finnish European allele frequency of variants with ≤ 15%WT activity (average allele frequency of 3.76 × 10^−5^ and those with > 15%WT (average allele frequency of 0:0014). The difference in the average allele frequency between the two groups was not statistically significant (p-value= 0:372, t-test).

We chose 15% WT as a threshold below which mutations are considered deleterious based on the fact that all but three out of 35 tested HGMD annotated pathogenic mutations reduced activity below this level, including three variants associated with attenuated disease (p.S612G, p.E634K, and p.L497V) [14–16]. The alleles associated with attenuated phenotype showed activity values in the range of 13% to 14% of WT. At this stage it is not clear why three of the putatively pathogenic HGMD mutations, p.A282V, p.P358L, and p.R482W, exhibited greater than 40% WT activity in our assay.

We evaluated the relationship between the enzymatic activity and allele frequency of tested variants (Figure 2 Insert B). We found that VUS with greater than 15% WT activity had higher allele frequencies, although this trend was not statistically significant. These data are consistent with the intuitive notion that high frequency variants are not likely to be associated with rare disease. We found that none of the tested variants with ≤ 15%WT activity had an allele frequency greater than 10^−3^.

### 2.3 Structural characteristics of functional variants

Using the publicly available structure of NAGLU (PDB ID 4XWH) we examined the structural characteristics of pathogenic variants (Section 4.4, Figure 3A). For this analysis we combined variants annotated as “DM” in HGMD with variants exhibiting ≤ 15%WT activity. When considering only mutations within the active-site pocket, almost all mutations (7 of 8) within within this region result in ≤ 15%WT activity. We found that, although most mutations in the active-site profoundly impacted activity, a variant’s lack of solvent exposure was more significantly associated with its impact on enzymatic activity than its distance from the active site. The observation has been reported previously [17, 18]

**Fig 3.**
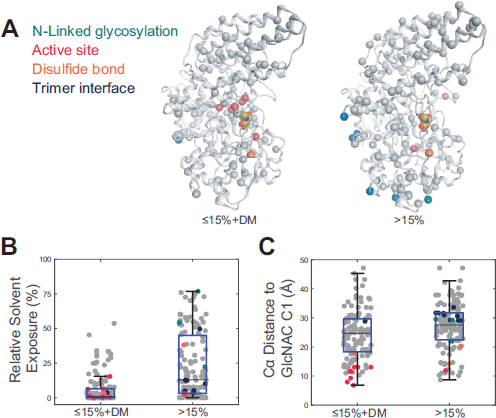
The structural characteristics of NAGLU variants. Mutations at N-linked glycosylation sites are shown in cyan, active-site pocket in red, disulfides in orange and trimer interface in blue. **(A)** Mutations (spheres) with ≤ 15%WT activity and DM variants in HGMD (left) and > 15%WT activity (right) mapped onto a NAGLU monomeric structure (PDB ID 4XWH) **(B)** The relative solvent exposure in the trimer NAGLU structure of variants with ≤ 15%WT activity and DM variants in HGMD compared to those with > 15%WT activity. ≤ 15%WT activity and DM variants were found to have an average percent solvent exposure of 5.96%. Variants with > 15%WT activity had an average percent solvent exposure of 23.20%. The differences in the average solvent exposure between the two groups was statistically significant at the 5% significance level with a p-value of 1.09 × 10^−12^. **(C)** The distance to the active site in the NAGLU structure of variants with ≤ 15%WT activity and DM variants in HGMD compared to those with > 15%WT activity. ≤ 15%WT activity and DM variants were found to have an average distance to the active site of 24.03Å. Variants with > 15%WT activity had an average distance to the active site of 27.23Å. The differences in the average distance to the active site between the two groups was statistically significant at the 5% significance level with a p-value of .001.

### 2.4 The statistical power of ExAC

We performed several calculations to explore whether current databases represent a large enough sample of the population in order to estimate the incidence of a rare disorder. For this calculation we used the beta distribution to calculate error margins and confidence intervals. We also assumed that only one fully penetrant variant contributes to disease. We found that ExAC, which at the time of data collection had the largest number of exomes among the publicly available databases, contains a sufficient number of samples in order to calculate disease incidence with a reasonable margin of error (Figure 4A). For the reported disease incidence of 1 in 321,128 for Sanfilippo Type B (9), the error margins with 95% confidence are between 1 in 428,070 (24% lower bound) to 1 in 249,879 (29% upper bound) given the overall ExAC sample size of 121, 412 (60, 706 individuals). The error margins increase to 33% and 39% for the lower and upper bounds respectively when limited to the Non-Finn European sample size of 66, 740 (33, 370 individuals). We believe these error margins are acceptable for the intended use of the dataset.

**Fig 4.**
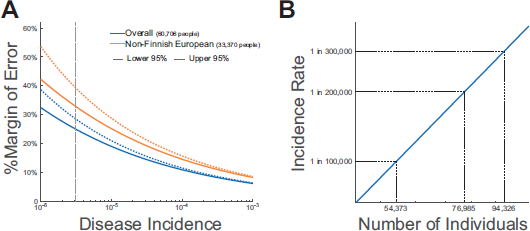
The Statistical Power of ExAC. **(A)** The percent margin of error the lower critical value would represent given a 95% confidence interval when calculating incidence using the same number of chromosomes in ExAC overall and for the Non-Finnish European cohort. The vertical dashed line represents the 1 in 312,128 Sanfilippo Type B incidence rate estimated [9]. **(B)** The number of individuals which should be sequenced such that the lower critical value will represent a 20% margin of error given a 95% confidence interval for a range of estimated incidence values.

We estimate that the overall and Non-Finnish European sample sizes of ExAC facilitate the calculation of disease incidence for disease as rare as 1 in 123,285 and 1 in 37,650 respectively with a 95% confidence interval and a 20% margin of error.

We also estimated the sample size necessary to achieve a 20% margin of error for a given disease incidence rate at a 95% confidence level (Section 4.11). As shown by Figure 4B, for a disorder as rare as 1 in 300,000, a sample size of roughly 95,000 individuals is necessary in order to obtain a 20% margin of error. While this is greater than the size of ExAC, which is made up of roughly 60,000 individuals, it is already exceeded by the number of individuals found in gnomAD.

In spite of having higher expected error margins, we used the Non-Finnish European sample in ExAC for all incidence calculations because it represents a more homogenous inter-breeding population than the overall ExAC population.

### 2.5 Sanfilippo Type B incidence estimates

We derived Sanfilippo Type B incidence estimates using the allele frequencies (Equation (3)) of three types of variants: 1.) HGMD variants 2.) LOF mutations, defined as nonsense, frameshift and splice site mutations not previously documented in HGMD, and 3.) VUS using a range of activity cut-offs (Figure 5A).

**Fig 5.**
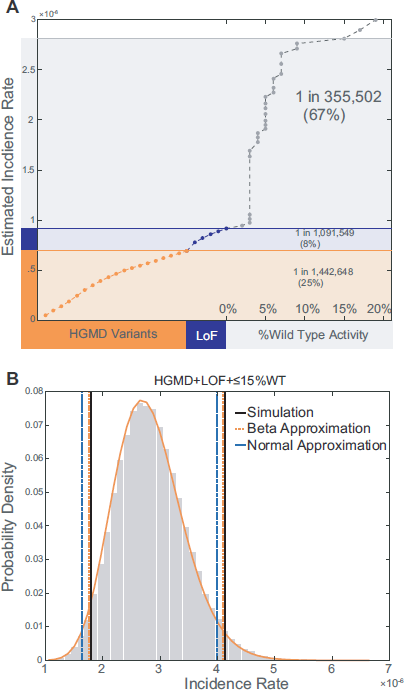
Allele frequency based Sanfilippo Type B incidence estimates. **(A)** The impact of each category of variants on our estimate of Sanfilippo Type B incidence. The contribution of HGMD variants, sorted by allele frequency, is shown in orange. The contribution of LoF variants which have yet to be documented in patients, also sorted by allele frequency, is shown in light blue. The contribution of VUS with %wt activity values below 15% is shown in grey. VUS are sorted by %wt values. Percentages in parenthesis represent the contribution of each category of variants to our final incidence estimate. **(B)** Confidence interval calculations. Grey bars represent the distribution of incidence rates observed through bootstrapping simulation. The orange solid line represents the distribution of incidence rates as modeled using the beta distribution. Using bootstrapping simulation we observed that 95% of simulated incidence values fell in the range of 1 in 558,306 and 1 in 241,749 for an equal tailed interval (vertical black lines). In comparison, using the beta-distribution we estimated that 95% of incidence values would fall in the range of 1 in 566,863 to 1 in 243,753 (vertical dashed orange lines). Using the normal approximation we estimated that 95% of incidence values would fall in the range of 1 in 610,093 to 1 in 250,830 (vertical dashed blue lines).

VUS with %WT values ≤ 15% had a profound impact on our estimate of disease incidence, and resulted in estimates that were close to values reported in the literature. Sanfilippo Type B incidence was estimated to be 1 in 355,502 when combining LoF and HGMD mutations with VUS with ≤ 15% WT activity. In contrast, the estimated incidence using only LoF and HGMD variants was 1 in 1,091,549.

Furthermore, as shown in Figure 5A, our incidence estimates were relatively insensitive to changes in the threshold used to define deleterious mutations from 10%WT to 15%WT activity. An increase in incidence of only 1.8% was observed when adding variants with 10 – 15% activity to our estimate.

### 2.6 Confidence interval calculations

Confidence interval estimates are impacted by sample size. Because sequence coverage varies across the genome, the actual sample size can also vary widely for each variant (S1 Fig). This variation for each allele in ExAC complicates the calculation of confidence intervals. We describe a statistical framework to address this complication (Section 4.6), and compare the resulting confidence intervals to what is observed through bootstrapping simulation (Section 4.9). Our framework facilitates the estimation of confidence intervals using either the normal approximation or the beta approximation of the binomial distribution.

The confidence intervals we calculated using the beta distribution produced more accurate estimates of the upper and lower bounds of the 95% confidence interval than the normal approximation, although both are in good agreement (Figure 5B). Using bootstrapping simulation we observed that 95% of simulated incidence values fell in the range of 1 in 558,306 and 1 in 241,749 for an equal tailed interval (vertical black lines) for our estimate of incidence using VUS with %WT values ≤ 15%. In comparison, using the beta-distribution we estimated that 95% of incidence values would fall in the range of 1 in 566,863 to 1 in 243,753 (vertical dashed orange lines). Using the normal approximation we estimated that 95% of incidence values would fall in the range of 1 in 610,093 to 1 in 250,830 (vertical dashed blue lines).

### 2.7 Using *in silico* predictors to estimate incidence

We first evaluated the agreement between *in silico* predictors of variant functional effect and the results of our activity assay. As shown in Figure 6A-B, PolyPhen and SIFT had roughly similar levels of sensitivity (or recall), but PolyPhen had higher specificity [19,20]. In spite of this, when we considered the quantitative scores of each predictor we observed that they were more correlated with each other than with our enzymatic activity values (S2 Fig).

**Fig. 6.**
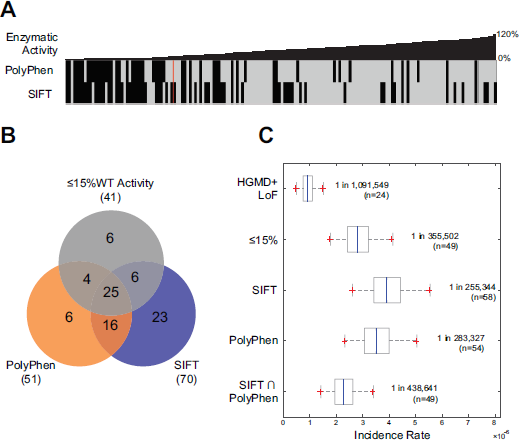
Performance of *in silico* predictors. **(A)** A comparison between the observed enzymatic activity for each variant, sorted from lowest to highest %WT activity, to the binary predictions from PolyPhen and SIFT, on the third row. A red vertical line represents the division between missense mutations with ≤ 15% WT activity and those with > 15% WT activity. **(B)** A Venn diagram showing the agreement between VUS observed to have ≤ 15% WT in our enzymatic activity assay and VUS categorized as “deleterious” by SIFT or “probably damaging” by PolyPhen. **(C)** Non-Finnish European incidence estimates obtained when considering only HGMD and LoF mutations (1 in 1,091,549), and when combining VUS with ≤ 15% WT activity (1 in 355,502), VUS categorized as “deleterious” by SIFT (1 in 255,344), VUS categorized as “probably damaging” by PolyPhen (1 in 288,327), or variants categorized as “deleterious” by SIFT and “probably damaging” by PolyPhen (1 in 438,641) with HGMD+LoF variants. VUS with an allele frequency in ExAC greater than 0.1% or for which one or more homozygous individuals were observed were excluded from estimates using SIFT and PolyPhen.

Selecting VUS using SIFT and PolyPhen produces reasonable incidence estimates but only after excluding variants with an allele frequency in ExAC greater than 0.1% or for which one or more homozygous individuals were observed. Combining those variants categorized as “deleterious” by SIFT or “probably damaging” by PolyPhen, with LOF and HGMD mutations produced incidence estimate of 1 in 255,344 and 1 in 283,327 respectively. Calculating incidence using the intersection of variants predicted by both tools, resulted in an estimated rate of 1 in 438,641. Without such filters, SIFT produced an incidence estimate of almost 100%. This was due to SIFT predicting two variants, R737G and R737S, as deleterious. These variants had a frequency of 92% and 1.9% respectively in the Non-Finnish European cohort of ExAC.

### 2.8 Comparison of results with Sanfilippo Type B rates reported in the literature

We compared our estimate of 1 in 355,502 for Sanfilippo Type B incidence to values reported in the literature (Table 1). Héron et al., estimated the overall incidence of Sanfilippo Type B in Europe to be 1 in 321,128. This estimate is only a 10% difference from what was calculated using the allele frequency of disease variants and knowledge about VUS. As they note, the incidence of Sanfilippo Type B has been documented in most European countries through retrospective epidemiological studies [9]. Incidence was reported to be the highest in Turkey (1 in 38,563), although it should be noted this rate was arrived at by considering Turkish individuals living in Germany [10]. At the other end of the spectrum, incidence was reported to be the lowest in Sweden (1 in 3,010,320) and the Czech Republic (1 in 4,261,897) [21, 22].

**Table 1.** A table of previously reported Sanfilippo Type B incidence rates.

| Publication | Date Range | Region/Ethnicity | Incidence |
| --- | --- | --- | --- |
| Baehner et al., 2005 [10] | 1980-1995 | Turkey | 1 in 38,563* |
| Al-Jasmi et al., 2013 [23] | 1995-2010 | UAE | 1 in 95,238 |
| Michelakakis et al., 1994 [24] | 1990-2006 | Greece | 1 in 122,423 |
| Pinto et al., 2004 [25] | 1966-1994 | Portugal | 1 in 138,888 |
| Meikle et al., 1999 [13] | 1980-1996 | Australia | 1 in 211,116 |
| Poorthuis et al., 1999 [26] | 1940-1991 | The Netherlands | 1 in 236,842 |
| Ben et al., 2009 [27] | 1970-2005 | Tunisia | 1 in 270,111 |
| Lin et al., 2009 [28] | 1984-2004 | Taiwan | 1 in 354,294 |
| Baehner et al., 2005 [10] | 1980-1995 | Germany | 1 in 387,694** |
| Heron et al., 2011 [9] | 1990-2006 | United Kingdom | 1 in 473,328 |
| Dionisi-Vici et al., 2002 [29] | 1985-1997 | Italy | 1 in 448,372 |
| Heron et al., 2011 [9] | 1985-1994 | France | 1 in 666,667 |
| Malm et al., 2008 [21] | 1975-2004 | Sweden | 1 in 3,010,320 |
| Poupetova et al., 2010 [22] | 1975-2008 | Czech Republic | 1 in 4,261,897 |

\*Estimated incidence in the Turkish population is based on the number of reported Sanfilippo Type B cases observed in Germany among births of individuals of Turkish ancestry (16 out of 617,013 births). \*\*Estimated incidence for Germany is adjusted to account for large fraction of observed individuals with Sanfilippo Type B being of Turkish ancestry. Before this adjustment the estimated incidence from [10] is 1 in 273,692.

## 3 Discussion

The data in this paper shows that ExAC enables reasonable estimates of disease incidence for rare monogenic disorders in the European population, as long as one takes into account the contribution of potentially pathogenic VUS. Our analysis illustrates this point; without considering the contribution of VUS, the estimated incidence of Sanfilippo Type B in Europe was 1 in 1,091,549. When including VUS with the potential to cause pathology, our estimate increased to 1 in 355,502, closely approximating the value reported in the literature of 1 in 321,128 [9]. Considering the large number of VUS in ExAC present in other genes (Figure 1), we anticipate this finding will be replicated. Thus, VUS should be considered when performing the same task for other disorders.

For many diseases, though, it might be difficult, expensive, or not feasible to develop an assay in order to test the impact of VUS on the activity of a particular gene product. In this case, one may improve the accuracy of incidence predictions by using *in silico* methods such as SIFT or PolyPhen. These methods are not reliable enough to predict the pathogenicity of individual variants and therefore should not be used to make clinical interpretations of patient genotypes on their own [30]. However, when the objective is to estimate the contribution to incidence of VUS, incorrect predictions for a small number of variants can be tolerated. We found that using PolyPhen and SIFT, alone or in combination, improved our estimate of disease incidence compared to ignoring the contribution of VUS altogether. These results were only arrived at by applying an allele frequency filter of 0.1% and excluding variants which occurred in the homozygous state in ExAC. Given that information regarding variant impact on NAGLU enzymatic activity was not used to train *in silico* predictors, our resulting set of 164 screened missense mutations also provide a unique data-set for the development and evaluation of new *in silico* methods [31].

We recognize that there are complexities to calculating disease incidence due to the consequence of certain allele combinations. Many pathogenic mutations that have high residual activity only cause disease when in the compound heterozygous state with a severe disease variant. This is the case with a very common splice site mutation, c.-32-13T>G, in Acid Alpha Glucosidase (GAA) which is associated with late-onset Pompe disease. This variant has an allele frequency of 3.5 × 10^−3^ in ExAC, but is rarely observed in the homozygous state in Pompe patients [32]. On the other hand, some common severe alleles are only found in the compound heterozygous state with a milder allele. This is the case with the Congenital Disorder of Glycosylation-1a (CDG-1a), caused by deficiencies in Phosphomannomutase 2 (PMM2). There have been no patients reported to be homozygous for the most common severe pathogenic variant (p.R141H), although the variant has frequently been reported as a compound heterozygote in patients, presumably due to prenatal lethality of the p.R141H/p.R141H genotype [33]. It is also expected that dominant or X-linked severe pathogenic alleles will be depleted in databases of presumably healthy individuals such as ExAC. Such exceptions need to be addressed when attempting to calculate disease incidence.

We have provided a theoretical framework in Sections 4.5 and 4.6 for calculating disease incidence and associated error margins. This theoretical framework assumes a randomly mating population at equilibrium [6,7]. This assumption is certainly not true, which can lead to an underestimate of disease incidence, even when multiple variants contribute to disease [34]. As shown in Table 1, the reported incidence rate of Sanfilippo Type B in Europe varies by orders of magnitude between countries. Given the exponential growth of population genomics data, allele frequency calculations will become more precise and facilitate the calculation of incidence for rarer disorders. Improved coverage for specific ethno/geographic groups will also enable meaningful calculations for specific populations. Collecting more data, and data that represents the population in a more granular, and unbiased manner will facilitate better disease incidence estimates. In spite of the anticipated growth in genomic data, VUS will still represent a significant challenge for incidence calculations, a challenge that can be addressed through improvements in *in silico* methods and increases in the throughput of experimental approaches.

## 4 Data and Methods

### 4.1 Selection of NAGLU variants for testing

We selected all NAGLU missense VUS from ExAC (release 0.3.1) for testing [8]. We used HGMD (2016 v1) as a source of disease associated variants [35]. In ExAC there are 207 variants that cause missense substitutions in the protein product NP 000254.2, associated with transcript NM 000263.3. Of these variants 18 failed to pass ExAC’s quality filter, resulting in 189 missense variants which we considered. In HGMD there are 90 missense variants associated with NAGLU annotated as being disease causing (assigned the DM or DM? tag). Of these 90 variants 26 were observed in ExAC, although two variants were annotated as failing ExAC’s quality filter. The remaining set of 165 missense variants not found in HGMD, which also passed ExAC’s quality filter, comprised the final set of missense VUS tested. We were able to successfully assay all but one (V117A) of these mutations for activity (Annotated as “VUS” as their source in S1 Table).

We also tested 38 pathogenic missense variants, 23 of which were found in ExAC as well as 8 nonsense variants, 3 of which had previously been associated with disease. These variants are listed in S1 Table with “HGMD” or “LOF” as their source.

SIFT and PolyPhen annotations for these variants were obtained from the ExAC VCF file and were generated using VEP v81.

### 4.2 Assaying the enzymatic activity of variants

NAGLU enzymatic activity testing was performed by HD BioSciences in Shanghai, China. Reference and mutant NAGLU variants were generated by cloning gene cassettes (BioMarin, unpublished) into the expression vector pCMV6 and transiently transfecting the resultant plasmids in equal parts into human embryonic kidney cells (HEK293, ATCC) using lipofectamine (11668, Invitrogen). Mutagenesis was performed using the QuikChange II XL site-directed mutagenesis kit according to the manufacturer’s instructions (200521, Stratagene). Each mutation was confirmed by DNA sequencing. As stated before, NP 000254.2 was used as the reference sequence.

HEK293 cells transfected with NAGLU plasmids were washed, then lysed 72 hours post-transfection. Enzyme activity was determined by adding 75 *μ*l 5 mM 4-Methylumbelliferyl-N-acetyl-α-D-glucosaminide (i.e., 4MU-NaGlu Substrate, Cat# 474500, Calbiochem) in assay buffer to 10 μl of cell lysate for 1 hour at 37°C. The reaction was stopped by adding 200 μl of glycine/NaOH buffer per well and the results read using a fluorescent microplate reader at Ex355nm Em460nm. Cell lysate protein concentration was determined using a BCA protein assay kit. Enzyme activity levels were normalized to lysate protein concentrations. Background activity from mock transfected cells was subtracted from the result for each construct. Results were then reported as the percentage wild-type activity of each mutation (%WT) by dividing the protein-concentration-normalized mutant activity by the normalized wild-type activity and multiplying by 100. Each construct was tested in at least three independent transfections. All enzyme assays were performed in duplicate and averaged.

### 4.3 Selection of variants for incidence calculation

In addition to tested VUS’s with low %WT values we utilized previously disease associated variants (annotated as DM in HGMD) observed in ExAC to calculate disease incidence. 23 variants were missense, three were stop gained and one was a splice affecting variant. These variants are listed in S2 Table with “HGMD” annotated as their source. We also found 14 variants categorized loss of function mutations (LOF). Such variants included frame-shift, premature stop gained variants and splice site variants. p.Trp743Ter was excluded from this set given that it exhibited high %WT activity. These variants are listed in S2 Table with “LOF” annotated as their source.

### 4.4 Structural analysis of mutations

Unless noted otherwise, MOE software was used to perform all structural analyses and generate molecular graphics [36]. A human NAGLU crystallographic structure (PDB ID 4XWH) was used to identify mutation-site residues involved in N-glycosylation modification, disulfide formation, active-site pocket, and biological assembly. Active-site pocket residues, defined as being located within 4.5 angstroms of a bound active-site ligand, were identified by superimposing the human NAGLU model with a structure of homologous bacterial family 89 glycoside hydrolase (PDB ID 2VCA) complexed with a reaction product, beta-N-acetyl-D-glucosamine (GlcNAc) [37]. Distances from a C-alpha atom of each mutation-site residue to a C-1 atom of the modeled GlcNAc were calculated using a custom MOE svl script. The atomic coordinates of a biological NAGLU trimer generated using COOT were used to identify those residues involved in subunit-subunit interactions at the trimer interface, and to calculate the accessible surface area of a mutation-site residue, expressed as percent surface exposures compared to the ideal surface determined from a Gly-X-Gly peptide [38, 39].

### 4.5 Estimating incidence

We describe methodology for estimating disease incidence with associated confidence intervals. The method we propose can be applied to any monogenic autosomal recessive disorder where multiple, fully penetrant, pathogenic variants contribute to disease.

If a single variant *v* is observed *k* times in a sample of 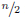 individuals from a diploid population then the frequency of that variant, *p*, is calculated as,

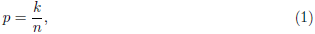

where *n* is the number of chromosomes.

In the simple, but very unlikely, case where a single variant causes disease the carrier rate is twice the allele frequency of the individual variant. If variants at more than one position in a gene contribute to disease we can calculate the probability at least one disease variant will appear at any position in a single copy of the gene. This is more easily calculated as one minus the probability no disease variant occurs,

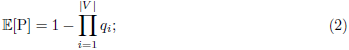

where *V* is the set of variants contributing to disease and *q_i_* is one minus *p_i_*, the observed frequency of disease variant 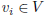. It should be pointed out that for very rare diseases calculating 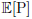 by taking the sum of all allele frequencies results in values that differ very little from what is arrived at using Equation (2).

We then estimated disease incidence 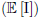 in a diploid population, assuming random mating and full penetrance of all variants, as the probability that a disease-causing variant occurs on both the paternal and maternal copy of the gene in an individual,

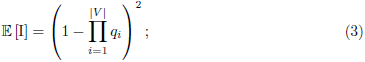

or more simply put as the square of the frequency of copies of the gene with at least one deleterious variant,

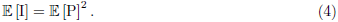

This allows us to calculate the expected incidence without enumerating over all 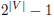 possible combinations of alleles.

### 4.6 Calculating the variance in incidence estimates

In order to calculate confidence intervals using the beta distribution or the normal approximation we first calculated the variance in our incidence estimate as,

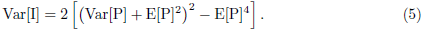

where E[P] is defined as above, and Var[P] represents the variance in the frequency of affected copies of the gene. Var[P] was calculated as the variance in the product of |*V*| random variables [40], as

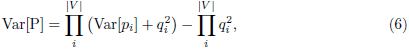

where Var[*p_i_*] is the variance in the frequency of variant 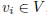, and takes the sample size associated with each individual variant into account.

For the purposes of this publication we modeled allele frequencies as a binomial distribution; calculating the variance of an individual variant’s frequency as,

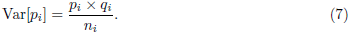

Both Equation (6) and Equation (5) remain the same when modeling allele frequencies using a hypergeometric distribution and calculating the variance in a single variant’s frequency as,

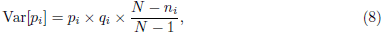

where *N* represents the overall size of the population being sampled.

When calculating 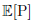 by taking the sum of all allele frequencies, Var[P] in Equation (5) can alternatively be calculated as,

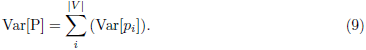

This methodology results in variance estimates that only differ slightly from what is generated using Equation (6), although the behavior of the two equations will be opposite when multiple variants contribute to disease. For example, holding sample size constant, a counter-intuitive observation is that the variance in incidence estimates using Equation (6) will be maximal when only one variant contributes to disease; as the number of variants increases, variance will decrease. Conversely, variance in incidence estimates using Equation (9) will be minimal when only one variant contributes to disease; as the number of variants increases, variance will increase. Variance in observed incidence will also be minimal using Equation (6) when the frequencies of the variants contributing to disease are uniform. This contrasts with the finding that the variance calculated using Equation (9) is the greatest when the probabilities of each random variable are uniform [41, Page 231].

### 4.7 Estimating confidence intervals using the beta distribution

Given the variance in our incidence estimate, Var[I], we calculated the *α* and *β* parameters for the beta distribution as,

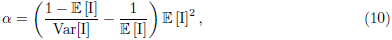

and,

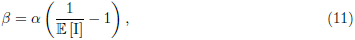

respectively.

We then used Newton’s method to find the critical 95% values of the inverse beta cumulative distribution given the *α* and *β* parameters.

### 4.8 Estimating confidence intervals using the normal approximation of the binomial distribution

Confidence intervals were also calculated using the normal approximation of the binomial distribution. Very simply, a two-tailed interval can be calculated as,

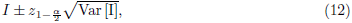

where 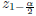 is the appropriate critical value for the desired percentage of coverage of the normal distribution.

### 4.9 Estimating error margins through bootstrapping simulation

We also calculated confidence intervals through bootstrapping simulation. This was carried out by re-estimating each variant’s allele frequency, which we will refer to as *p*’, by two methods independently; resampling with replacement and using a binomial random function. Parameters for the binomial random function were set such that the number of trials were equal to the sample size, *n*, associated with each variant, and the likelihood of success was set to be the frequency, *p*, of each variant respectively. This method is modifiable so that confidence intervals can be modeled using the hypergeometric distribution, or sampling without replacement. Once each variant’s frequency was re-estimated the disease incidence, which we will refer to as *I*’, was re-estimated using Equation (3). This process was carried out over 10^5^ iterations, generating a distribution of incidence estimates. Confidence intervals were then estimated by considering the value *I*’ for which the defined percentage of observed incidence values from the simulation are greater than or less than.

Numerical simulation was carried out in MATLAB. Allele frequencies were re-estimated using the bootstrp or binornd functions, and simulation was carried out using the parallel computing toolbox which relies on the combined multiple recursive random number generator. Code has been provided in the supplemental materials.

### 4.10 Using the average sample size to calculate confidence intervals

We also explored calculating confidence intervals using the sum of allele frequencies and the average sample size for all variants. In order to do this we modified Equation (7) to consider the sum of allele frequencies and their average sample size,

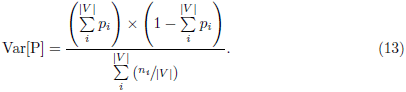

The resulting estimate of Var[P] was then used as input for Equations (5) and (12).

We found that this technique will only be accurate when there is very little variation in the sample size associated with each variant. Using the average sample size it was estimated that 95% of incidence values would fall in the range of 1 in 587,202 to 1 in 254,337 for the same data-set as above. While these values are not a drastic departure from what was observed through bootstrapping simulation, there was very little variation in the sample size associated with these variants in the Non-Finnish European population in ExAC (S1 Fig).

Changing the sample size associated with just a single variant, Ser612Gly, from 40,638 to 10,000, resulted in a more drastic departure between this method and what was observed in simulation (S3 Fig). With this modification we observed that 95% of bootstrap simulated incidence values fell in the range of 1 in 629,241 to 1 in 226,602 (vertical black lines). While the beta-distribution estimated that 95% of incidence values would fall in the range of 1 in 643,633 to 1 in 225,238 (vertical dashed orange lines), using summed allele frequencies and the average sample size produced a range of 1 in 589,282 to 1 in 253,949 (vertical dashed cyan lines). Data-sets with more variation in sample size will generate even larger departures between this method and what will be observed through bootstrapping.

### 4.11 Calculating power of current data-sets

In some cases it may be necessary to determine the number of individuals that must be sequenced in order to estimate the incidence of a disorder of a particular rarity within an acceptable margin of error. For the beta distribution this was done by iterating through values of *n* in order to obtain the appropriate percent margin of error for a given incidence rate. As it may be the case no single value of *n* will produce an exact desired margin of error value, we chose the *n* with the closest margin of error.

We also provide formulas for such estimates using the normal approximation of the binomial distribution. Given a single variant with allele frequency *p* contributes to disease, the variance in the expected incidence can be written as,

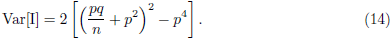

Solving Equation (12) for *n* using Equation (14) one obtains the form,

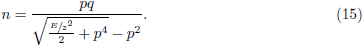

We use Equation (15) to calculate the number of chromosomes that must be sequenced in order to obtain a given margin of error and confidence interval. Note that, as humans are diploid, 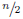 represents the number of individuals necessary to obtain a particular margin of error.

## Supporting information

**S1 Fig.**
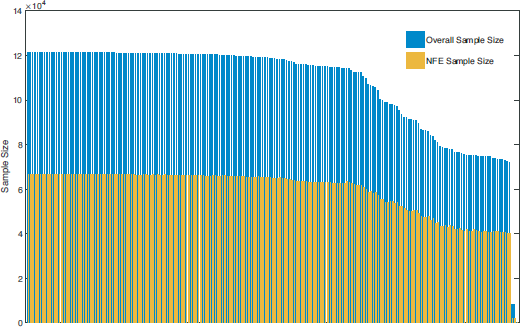
Variant Sample Sizes. The Overall and Non-Finnish European sample size in ExAC associated with each mutation used in the HGMD+LOF+≤ 15% incidence calculation. p.Glu284GlyfsTer33 and p.Leu281Pro had the largest associated Non-Finnish European sample size of 66,740. In contrast, c.383+1G>T had the smallest associated NonFinnish European sample size of 1,962. This variant did not contribute to our Non-Finnish European incidence calculation as it had an allele frequency of 0 in this population.

**S2 Fig.**
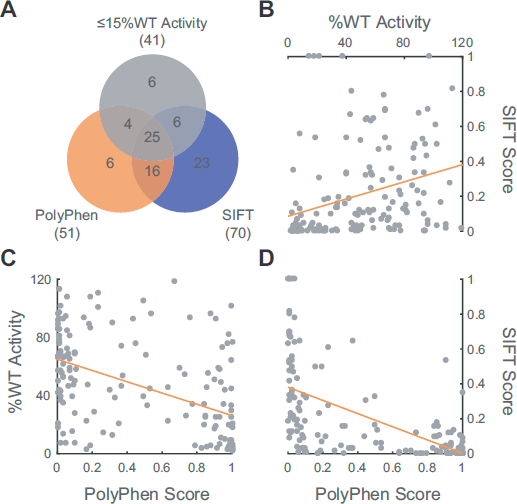
Correlation of SIFT, PolyPhen and observed activity values. **(A)** A Venn diagram showing the agreement between VUS observed to have ≤ 15% WT in our enzymatic activity assay and VUS categorized as “deleterious” by SIFT or “probably damaging” by PolyPhen. **(B)** A scatterplot of observed %WT activity values and SIFT scores. A correlation coefficient of 0.3147 was observed between %WT activity values and SIFT scores. **(C)** A scatterplot of observed %WT activity values and PolyPhen scores. A correlation coefficient of −0.4941 was observed between %WT activity values and PolyPhen scores. **(D)** A scatterplot of observed SIFT and PolyPhen scores. A correlation coefficient of −0.6159 was observed between SIFT and PolyPhen scores.

**S3 Fig.**
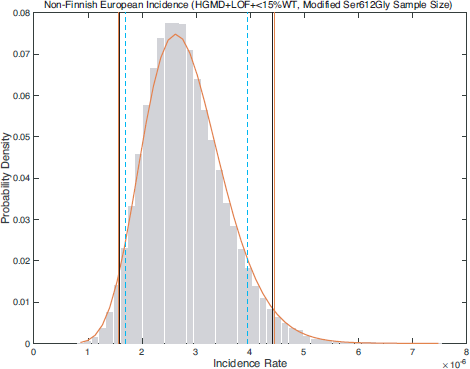
Modified sample size confidence intervals. Confidence intervals observed when changing the sample size associated with Ser612Gly from 40,638 to 10,000. Grey bars represent the distribution of incidence rates observed through bootstrapping simulation. The orange solid line represents the distribution of incidence rates as modeled using the beta distribution. Using bootstrapping simulation we observed that 95% of simulated incidence values fell in the range of 1 in 629,241 to 1 in 226,602 for an equal tailed interval (vertical black lines). Using the beta-distribution we estimated that 95% of incidence values would fall in the range of 1 in 643,633 to 1 in 225,238 (vertical dashed orange lines). Using summed allele frequencies and the average sample size produced a range of 1 in 589,282 to 1 in 253,949 (vertical dashed cyan lines).

**S1 Table. Tested Variants.** A table of tested NAGLU variants and their observed %WT activity values.

**S2 Table. Allele frequencies.** A table of the variant allele frequencies and sample sizes from ExAC. When available, %WT activity values are also included.

## Acknowledgements

We would like to acknowledge Josh Woloszynek, Yuanbin Ru, Predrag Radivojac, and Yaniv Erlich for their comments on the manuscript. We would also like to thank Larry Hengl for his assistance with database management.

